# Inflammatory stress-mediated chromatin changes underlie dysfunction in endothelial cells

**DOI:** 10.1101/2023.10.11.561959

**Authors:** Haibo Liu, Amada D. Caliz, Heather Learnard, Milka Koupenova, John F. Keaney, Shashi Kant, Lihua Julie Zhu, Anastassiia Vertii

## Abstract

Inflammatory stresses underlie endothelial dysfunction and contribute to the development of chronic cardiovascular disorders such as atherosclerosis and vascular fibrosis. The initial transcriptional response of endothelial cells to pro-inflammatory cytokines such as TNF-alpha is well established. However, very few studies uncover the effects of inflammatory stresses on chromatin architecture. We used integrative analysis of ATAC-seq and RNA-seq data to investigate chromatin alterations in human endothelial cells in response to TNF-alpha and febrile-range heat stress exposure. Multi-omics data analysis suggests a correlation between the transcription of stress-related genes and endothelial dysfunction drivers with chromatin regions exhibiting differential accessibility. Moreover, microscopy identified the dynamics in the nuclear organization, specifically, the changes in a subset of heterochromatic nucleoli-associated chromatin domains, the centromeres. Upon inflammatory stress exposure, the centromeres decreased association with nucleoli in a p38-dependent manner and increased the number of transcripts from pericentromeric regions. Overall, we provide two lines of evidence that suggest chromatin alterations in vascular endothelial cells during inflammatory stresses.

## INTRODUCTION

Inflammation is an evolutionary conserved 600 million-year-old biological response that is instrumental for the immune response during infections ^1,2^. Pro-inflammatory cytokines mediate the immune response against viral and bacterial infections. However, prolonged exposure to pro-inflammatory cytokines contributes to various human pathologies, including cardiovascular diseases such as atherosclerosis and vascular fibrosis ^3^. Atherosclerosis, the leading cause of myocardial infarction, stroke, and peripheral vascular disease, is a multi-stage disease that initially involves dysfunctional and inflamed endothelium, which triggers an inflammatory response leading to the recruitment of inflammatory cells, including monocytes, T-cells, and smooth muscle cells de-differentiation ^4^. Endothelial dysfunction is defined as a shift in cellular metabolism towards an inflammatory response, including cell surface profile alterations and proliferative changes ^3^. In contrast, a healthy inner vascular lining consists of non-migratory endothelial cells with a cell surface molecular profile that prevents the recruitment of pro-inflammatory factors, thereby preventing the initiation of atherosclerosis and many more inflammatory vascular diseases ^5^. Exposure of vascular endothelial cells to pro-inflammatory cytokines, such as tumor necrosis factor-alpha (TNF-alpha), induces the expression of cell adhesion molecules (CAMs), such as vascular cell adhesion molecule-1(VCAM-1) and intercellular adhesion molecule-1(ICAM-1), which are necessary for leukocyte recruitment, thus initiating endothelial dysfunction ^4–8^. Consequently, endothelial cells undergo mesenchymal transition (EndMT), thus changing their cell fate. The EndMT process is often considered as a type of epithelial-to-mesenchymal transition (EMT) ^9^. EndMT is a prerequisite for vascular fibrosis, a hallmark of multiple immune-related diseases ^3^.

At the cellular level, inflammation can manifest through exposure to pro-inflammatory cytokines such as TNF-alpha and febrile-range heat stress. Exposure to TNF-alpha leads to cell-type-dependent changes in patients with chronic inflammation ^10^. For example, in rheumatoid arthritis patients, the non-immune cells responsible for removing debris or synoviocytes, but not macrophages, demonstrated prolonged expression of genes with a single TNF-alpha pulse accompanied by an increase in chromatin accessibility ^10^. Endothelial responses to inflammatory stress largely rely on the mitogen-activated kinase (MAPK) signal transduction, specifically, p38MAP kinase (p38MAPK) and c-Jun N-terminal kinase (JNK), both well-known stress-activated protein kinases, due to their essential roles in orchestrated stress responses, including cytokines ^11,12^. Despite the significant role of inflammation in vascular disorders, the consequences of inflammatory stimuli on the chromatin organization of vascular endothelial cells and the contribution of febrile-like fever to endothelial dysfunction are poorly understood.

Centromeres are heterochromatin regions critical for establishing the kinetochores during mitosis, featuring centromere-specific histone CenpA embedded in the repetitive DNA, termed alpha-satellite repeats ^13^. It is well-known that in interphase nuclei, the centromeres localize close to nucleoli, representing one of the types of nucleoli-associated chromatin domains, NADs ^14,15^. The dynamics between centromere-nucleoli association is known to change during cell differentiation and cancer ^16^. Disruption of centromere-nucleoli association leads to an increase in alpha-satellite transcription ^17^.

In this study, our objective is to explore the impact of inflammatory stresses induced by either heat stress or cytokine exposure on chromatin accessibility, organization, and transcriptional responses. Additionally, we investigated the potential impact of TNF-alpha exposure and febrile-like heat stress on the centromeres - nucleoli association.

## RESULTS

### Short-term exposure to inflammatory stresses changes chromatin accessibility in a stress-type-dependent context

Fever is an essential component of inflammation. However, how febrile-like temperature stress (heat stress, HS) affects chromatin dynamics in endothelial cells is unknown. Here, we report that human endothelial cells subjected to short-term (two to six hours) inflammatory stresses alter chromatin accessibility and transcriptional response. To mimic the fever scenario, we used febrile-range heat stress of 39-40ºC for two hours ^18^. To investigate genome-wide changes in chromatin accessibility, we performed ATAC-seq on EC cells, exposed to either 2hr of 39ºC febrile-like heat stress or 6hr of 10 ng/ml TNF-alpha (Fig. 1 and Fig. S1). Our biological replicates were similar to each other with consistently less reads in heat-stressed cells (Fig. S1A, B, C, D). Most ATAC-seq peaks were located within distal intergenic and less than one kb promotor regions (Fig. 1A). Out of 79132 peaks, we observed 18494 differentially accessible regions (DARs) following heat stress as compared to 6023 after TNF-alpha treatment (Fig.1 B, C), suggesting HS had a greater effect on chromatin accessibility than the pro-inflammatory cytokine TNF-alpha. The ATAC-seq motif deviation analysis revealed that TNF-alpha-induced accessible regions contain multiple binding sites of classic cytokine-mediated sets of transcription factors such as RelA, NFkB, and STAT1 (Fig. 1D). Similarly, HS-induced accessible regions contain binding sites of classic heat shock transcription factors (TFs) including HSF1, HSF2, HSF4, and downstream of MAPKs TFs FOS, and Jun (Fig. 1D). Such stress-specific responses are also observed in KEGG pathway overrepresentation analysis (Fig.1 E and F, respectively). For example, TNF-specific response and cytokine-cytokine pathways are enriched in TNF-treated samples. Meanwhile, there are common enriched pathways in both heat stress and cytokine exposure, such as cancer pathways and lipid and atherosclerosis pathways, which are indicative of inflammation-induced changes in endothelial cells and the initiation of EC dysfunction. Taken together, temperature stress and pro-inflammatory cytokine exposure lead to changes in chromatin accessibility in HUVEC cells, and these changes involve stress-specific responses as well as overlapping pathways that may contribute to EC dysfunction.

**Figure 1.**
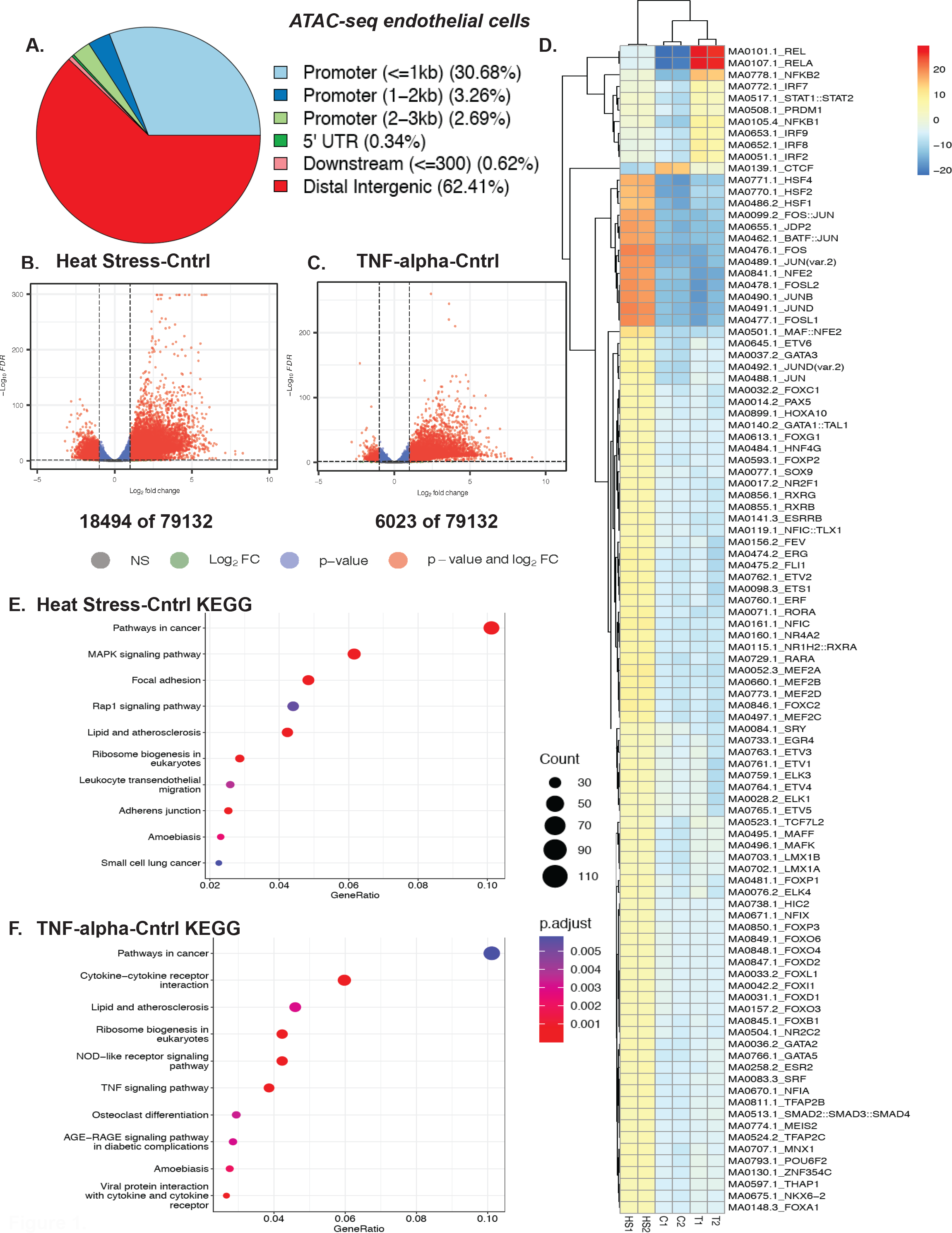
Chromatin accessibility assessment of febrile-like HS and TNF-alpha-treated endothelial cells revealed stress response and endothelial dysfunction drivers. A. Distribution of peaks among genomic features. B. A volcano plot of HS-Cntrl differentially accessible regions (DARs) ATAC-seq regions. C. A volcano plot of TNF-Cntrl (DARs) ATAC-seq regions. D. ATAC-seq motif deviation from HUVEC cells treated with febrile-like HS (39°C for 2hrs) or TNF-alpha (10ng/ml for 6hrs) (C1, C2, T1, T2, and HS1, HS2 are biological replicates of control, TNF-alpha, and heat stress treated cells). E. HS-Cntrl up-regulated KEGG pathway overrepresentation analysis. F. TNF-Cntrl up-regulated KEGG pathway overrepresentation analysis.

### Febrile-like HS and TNF-alpha induce a profound transcriptional response, including stress-specific EC dysfunction drivers

Differences in chromatin accessibility often underlie changes in gene regulation. Transcriptional response to inflammatory stresses was assessed using RNA-seq with exactly the same cell treatments as for ATAC-seq (Fig. 2 and Fig.2S). As expected, the two stresses lead to profound changes in gene expression (Fig. 2 A, B, C). Notably, HS and TNF-alpha induced the expression of cell adhesion molecules (CAMs) that are instrumental in the adhesion of blood monocytes and other leukocytes to EC cells. Upregulation of CAMs and genes mediating EndMT is often considered a prerequisite for EC dysfunction. Our results suggest both the HS and TNF-alpha induced upregulation of EC dysfunction drivers (Fig. 2 D and E, respectively). Although EC under HS and TNF-alpha exposure expressed dysfunction markers, the specific genes often differ between the two stresses (Fig. 2 F). For example, HS but not TNF-alpha-induced PECAM1 (CD31), an adhesive molecule essential for leukocyte migration through the EC layer ^19^. Conversely, TNF-alpha but not HS-induced ICAM1, a cell surface glycoprotein that is also instrumental in leukocyte recruitment to EC cells during inflammation ^20^. Of interest, both stresses induced KLF4, a multifunctional transcription factor, critical for inflammation resolution in EC cells.

**Figure 2.**
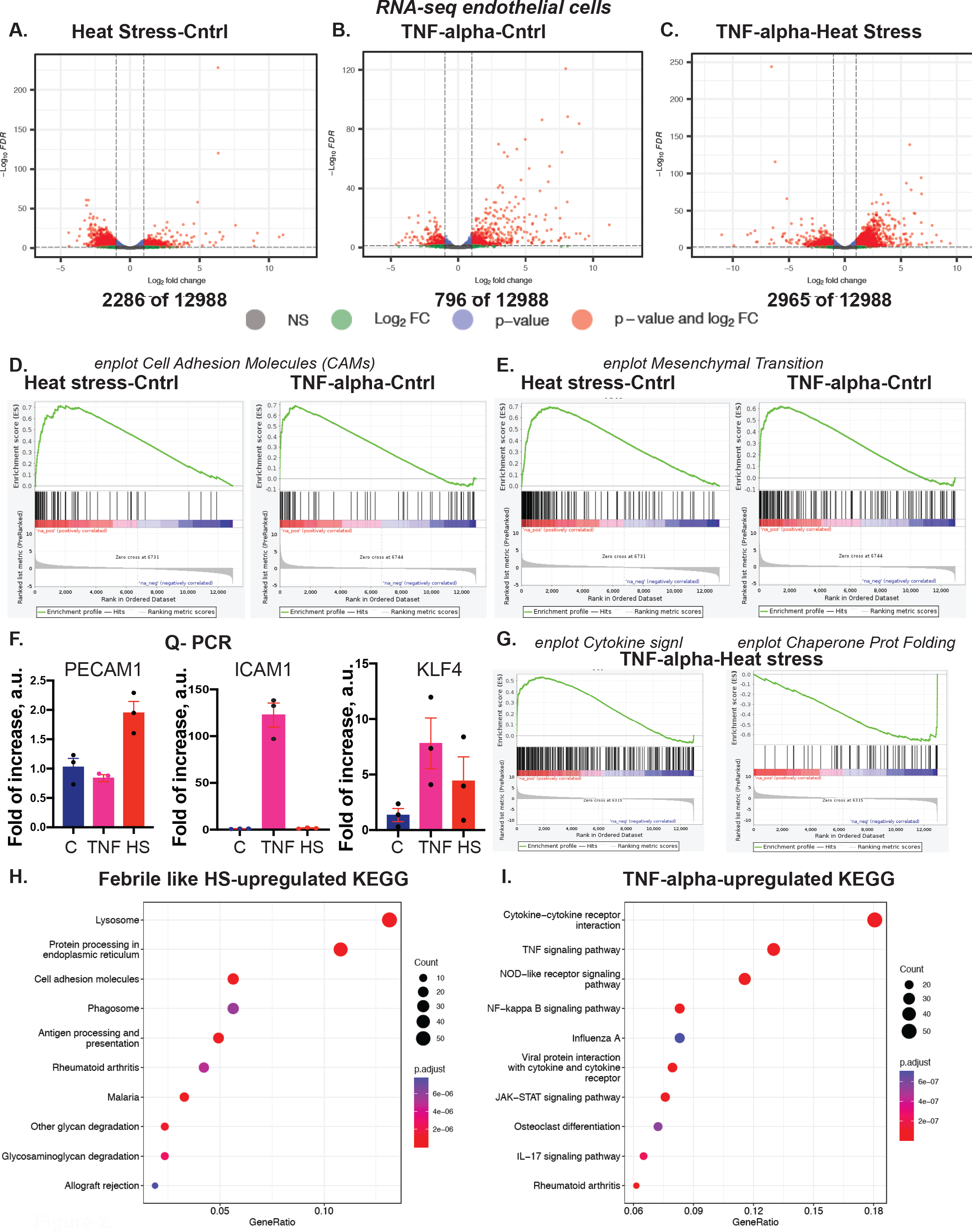
Febrile-like HS and TNF-alpha induce a profound transcriptional response, including EC dysfunction drivers. A. A volcano plot of HS-Cntrl Differentially Expressed Genes, RNA-seq. B. A volcano plot of TNF-Cntrl Differentially Expressed Genes, RNA-seq. C. A volcano plot of TNF-HS Differentially Expressed Genes, RNA-seq. D. Gene Set Enrichment Analysis (GSEA) enrichment plot for cell adhesive molecules, CAMs. E. GSEA enrichment plot for mesenchymal transition genes. F. Q-PCR to confirm stress-induced upregulation of markers for EC dysfunction, CAMs, and anti-inflammatory protein KLF4. G. GSEA enrichment plots for cytokine signaling and chaperone-mediated protein folding for TNF-HS. H. KEGG analysis for HS-Cntrl and I. for TNF-Cntrl RNA-seq samples.

Similar to ATAC-seq, the two stresses induced stress-specific genes, such as cytokine signaling that is specific for TNF-alpha and chaperone-mediated protein folding genes, characteristic of HS (Fig. 2 G). Additionally, KEGG pathway analysis for HS and TNF-alpha reveals inflammatory yet specific for each stress pathway genes (Fig. 2 H and I). The stress-specific gene set contains well-characterized transcriptional responses and the induction of EC dysfunctional genes that differ between the two stresses.

### Integrative analysis of ATAC-seq and RNA-seq revealed stress-specific responses in EC dysfunction genes

To assess whether changes observed in chromatin accessibility underlie transcriptional response, we performed integrative analysis of ATAC-seq and RNA-seq data from HUVEC cells. To visualize the integrative analysis results and determine genes whose transcriptional activation is accompanied by an increase in chromatin accessibility, we plotted log2 DEG (differentially expressed genes) changes vs. log2 DAR (differentially accessible regions) changes for HS vs. control and TNF-alpha vs. control (Fig. 3A and B, Fig. S3 A and B, respectively). The increase in chromatin accessibility often correlated with an increase in stress-specific and EC dysfunctional gene expression (Fig. 3A and B). For example, HS induced both DARs and DEGs in heat shock proteins, HSPs (stress-specific) genes, PECAM1, and HDAC9 (EC dysfunctional). Similarly, TNF-alpha induced both chromatin and transcriptional changes in SELE, TNF, CPEB4, and others. One of the critical questions in the field is whether the 3D chromatin organization changes during inflammation. ATAC-seq informs about the chromatin accessibility, which can be affected by multiple scenarios, including alterations in transcription factor binding and the organization of densely packaged heterochromatin domains ^21^. We next ask whether the DARs in EC cells during inflammatory exposure were found in heterochromatin or euchromatin regions. For this purpose, we used Hi-C data from untreated EC cells (ENCODE data accession number: ENCFF005ZBU) to visualize heterochromatin and euchromatin domains (gene silenced compartment B (orange) and gene active compartment A (magenta) in Hi-C data, respectively). Interestingly, DARs in genes of interest were found in both compartments. Specifically, VCAM1 DARs were detected in compartment A, while VCAN DARs were detected in compartment B (Fig. 3C and D, respectively). VCAN encodes for versican protein, a large protein that contributes to extracellular matrix formation and is expressed by inflamed endothelium ^22^. Of particular interest is the observation that in some chromosomes, DARs were more likely to border compartment B, and in other chromosomes, the changes were distributed rather uniformly across the chromosome. For example, in chromosome 9 (chr.9), DARs from HS and TNF samples were found across the whole chromosome. However, in chromosome 11 (chr.11), DARs are less likely to be found in heterochromatic compartment B (Fig. 3E). In sum, we identified DARs that are underlying gene expression responses in EC cells exposed to short-term inflammatory stresses. We also observed a stress-specific set of DARs/DEGs, but whether they have a cumulative effect during inflammation remains unknown.

**Figure 3.**
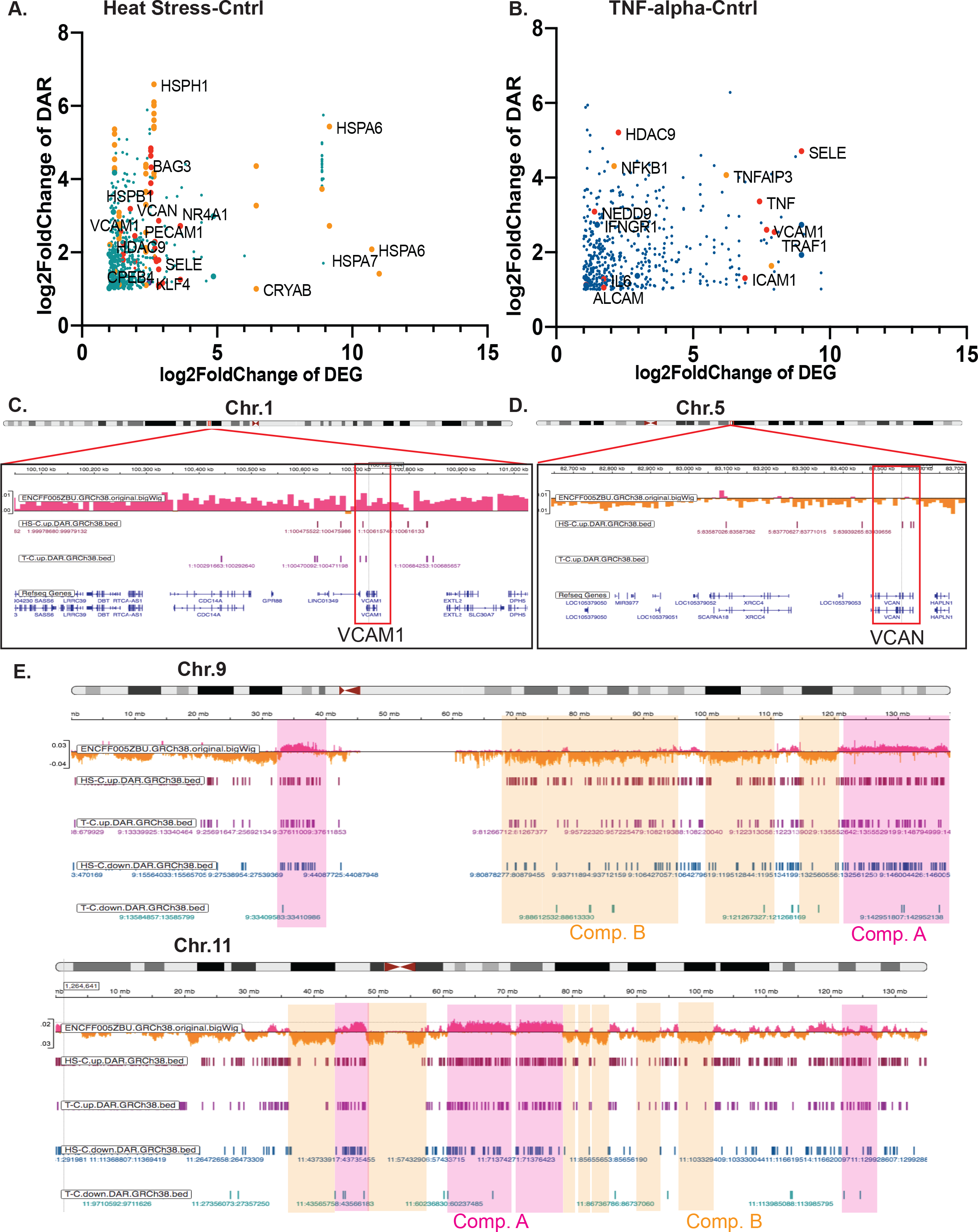
Transcriptional changes in multiple EC dysfunction markers also undergo alterations at the chromatin accessibility level. A. Integrative analysis of ATAC-seq and RNA-seq for HS-Cntrl and B. for TNF-Cntrl (in orange color are examples of stress-specific genes, in red are examples of EC dysfunctional genes). C. An IGV browser snapshot of a region of chr.1 with human endothelial vein cells Hi-C track (ENCODE data accession number: ENCFF005ZBU), HS-T upregulated DARs, and TNF-C upregulated DARs at the site of VCAM1 gene. D. An IGV browser snapshot of a region of chr.5 with human endothelial vein cells Hi-C track (ENCODE data accession number: ENCFF005ZBU), HS-T upregulated DARs track and TNF-C upregulated DARs track at the site of VCAN gene. E. An IGV browser snapshot of a chr.9 and chr.11 with human endothelial vein cells Hi-C track (ENCODE data accession number: ENCFF005ZBU), HS-T upregulated and downregulated DARs tracks, and TNF-C upregulated and downregulated DARs tracks suggests more ATAC-seq DARs are within compartment A than compartment B.

### Inflammatory stresses decrease centromere-nucleoli association in a p38 MAPK-dependent manner

In interphase nuclei, the centromeres typically localize in close proximity to nucleoli and represent a subset of NADs. In our ATAC seq analysis, we identified chromatin changes in a number of centromere-related genes, specifically, centromere-specific histone CENPA, an alpha-satellite-repeat binding centromere protein CENPB, and centromere protein CENPC which is critical for centromere-nucleoli association (Fig. 4A). Integrative analysis of ATAC-seq and RNA-seq analysis of febrile-like heat stress samples vs. control revealed an upregulation of otherwise developmentally regulated transcription factor gene ZNF423 (Chr.16: 49487524..494859279), located within compartment B in the close proximity to pericentromeric region (Fig. 4A). Zinc finger (ZNF) clusters often localize around nucleoli in interphase cells and represent gene-silenced nucleoli-associated heterochromatic chromatin domains, NADs ^23^. We, therefore, investigated NADs-nucleoli dynamics during HAEC and HUVEC exposure to inflammatory stresses using confocal microscopy. First, we confirmed the inflammatory response in HAEC cells, which includes the expression of CAM molecules, which are characteristic of the initial stage of TNF-alpha-induced EC dysfunction. Therefore, we assessed the upregulation of genes such as vascular cell adhesion molecule-1(VCAM-1), intercellular adhesion molecule-1(ICAM-1), and pro-inflammatory cytokine IL6 in HAEC cells (Fig. 4E). We next determined nucleoli-centromere dynamics. For this purpose, we used a well-established centromere marker CREST to visualize centromeric NADs and nucleolin antibodies to visualize nucleoli (Fig. 4B, C, F, G). We observed an increased dissociation of centromeres from nucleoli in TNF-alpha-challenged cells (Fig. 4B and C). Centromere-nucleoli association is essential for the regulation of centromere transcripts, and the depletion of key regulators of this association, such as CENPC, leads to an increase in pericentromeric alpha-satellite transcripts (ASAT) ^17^. We therefore tested whether TNF-alpha-induced dissociation changes alpha-satellite transcripts using the same as previously published RNA-fluorescence in situ hybridization (FISH) probe (Fig. 4D) ^17^. Our data suggest that TNF-alpha exposure induces ASAT transcripts as a consequence of nucleoli-centromere dissociation. We next confirmed the TNF-induced decrease in nucleoli-centromere association in HUVEC cells and tested the contribution of HS to nucleoli-centromere dynamics. HS decreased centromere-nucleoli association, similar to TNF-alpha-treated cells (Fig. 4F and G). MAPK p38 has pleiotropic effects on cellular metabolism and is instrumental during inflammatory response ^24^. We probed for the contribution of p38 activation on nucleoli-centromere dynamics during TNF-alpha and HS exposure (Fig. 4F and G). For this purpose, we used well-known small molecule inhibitors of p38 kinase activity, BIRB796 and SB203580, currently in clinical trials ^25^. Our data show that inhibition of p38 kinase activity restored centromere-nucleoli association, suggesting that p38 MAPK mediates the nucleoli-centromere dynamics during inflammatory stresses (Fig. 4F, G, H).

**Figure 4.**
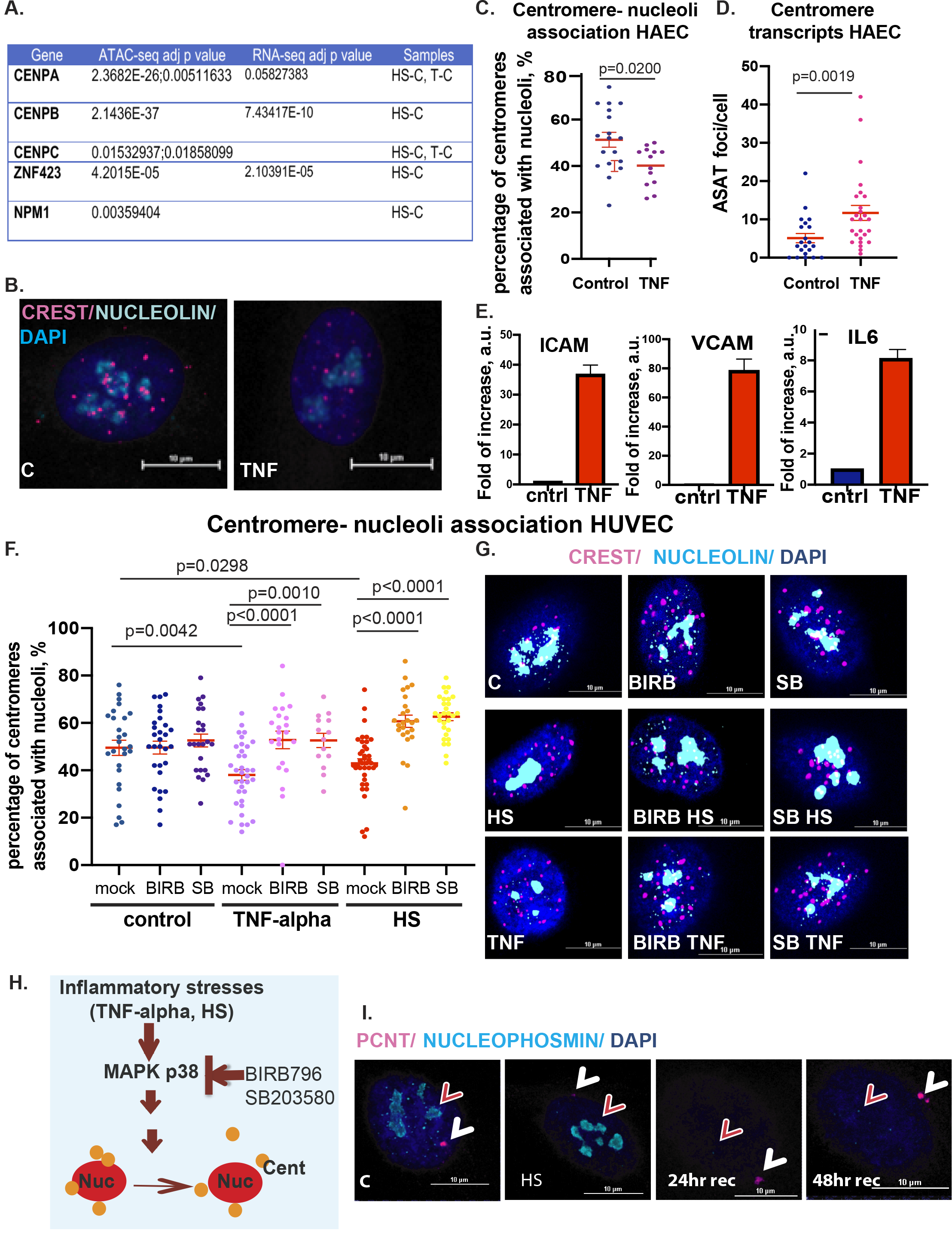
Dynamics in centromere-nucleoli association during inflammatory stress exposure depends on p38 MAPK kinase. A. Changes in centromere genes chromatin accessibility during exposure to inflammatory stresses. B. Confocal microscopy images of the centromere (CREST, magenta) -nucleoli (nucleoli, cyan) association in control and TNF-alpha treated HAEC cells (scale bar 10 μm). C. Quantification of nucleoli-centromere association: each dot represents the percentage of nucleoli-associated centromeres per nuclei. D. Quantification of alpha satellite (ASAT) transcripts per cell in control and TNF-alpha-treated cells. E. Q-PCR for endothelial dysfunction genes ICAM1, VCAM1, and IL-6 upregulation as a result of TNF-alpha exposure in HAEC cells. F. Quantification of nucleoli-centromere association in TNF-alpha and HS-treated HUVEC cells; each dot represents the percentage of nucleoli-associated centromeres per nuclei. When indicated, cells were exposed to stresses in the presence of p38MAPK inhibitors BIRB and SB. G. Confocal microscopy images of the centromere (CREST, magenta)-nucleoli (nucleoli, cyan) association in control and TNF-alpha treated HUVEC cells in the presence of p38 inhibitors, BIRB and SB (scale bar 10 μm). H. Model of the role of p38 in centromere-nucleoli dynamics during inflammatory stress. I. Nucleoli marker nucleophosmin (cyan, red arrow) disruption during HS recovery. Centrosome (pericentrin marker, PCNT, magenta, white arrow) is used as an intracellular marker for cellular recovery from stress in HUVEC cells, scale bar 10 μm.

Several molecules were previously reported to contribute to the nucleoli-centromere association, such as nucleolin interactions ^26,27^, fibrillarin ^17^, Ki67 ^28^, chromatin assembly factor CAF1 ^29^, CenpA ^17^, and H3K27me3 ^23^. One of the major interacting partners of CenpA is nucleoli protein nucleophosmin (NPM1) ^30^, which came as a significantly changed chromatin accessibility gene in ATAC-seq data from heat stress samples (Fig. 4A). Nucleophosmin decorates the outermost layer of the tri-partite nucleoli structure that faces nucleoli-associated chromatin domains. Therefore, we tested the nucleophosmin localization during inflammatory stresses and recovery after febrile-like heat stress (Fig. 4I). The centrosome and centromeres are critical for establishing mitotic machinery and proper separation of genetic material into two daughter cells. Centrosome is an established stress-sensing organelle that undergoes alteration during heat stress and returns to normal within 14-24 hr following the treatment ^31^. We monitored its *bona fide* protein, pericentrin (PCNT, magenta), as an intracellular marker for cell stress recovery dynamics. Indeed, the pericentrin signal was decreased during stress and recovered after 24 hours of returning to normal temperature (Fig. 4I). In contrast to PCNT, we did not observe significant changes in NPM1 immediately after HS. Instead, the dramatic decrease in NPM1 signal occurred during the recovery period, starting at 24 hr. The dissociation of CenpA-containing centromeres from nucleoli began immediately after heat stress (Fig. 4F, G), suggesting that dissociation precedes changes in nucleoli integrity.

To summarize, inflammatory stress-mediated alteration of centromere-nucleoli association depends on p38 and correlates with transcriptional activation of satellite repeats. Our data outlines previously unreported dynamics in nucleoli protein nucleophosmin, distinct from another well-established stress-sensing organelle, the centrosome ^31–33^. This data suggests that the two stress-sensing organelles, the centrosome and nucleoli, have different timing in their recovery from febrile-like heat stress. As centrosome is essential for cell division, the question of whether the cells undergo cell divisions with recovered centrosomes yet with impaired nucleoli composition remains to be investigated.

Both heat stress and cytokine exposure lead to an increase in the expression of stress-specific EC dysfunction genes. This transcriptional response was accompanied by an increase in chromatin accessibility. Moreover, the inflammatory stresses affect chromatin organization at the level of nucleoli-associated chromatin domains (NADs), specifically, a subset of NADs, the centromeres. Here, we report that TNF-alpha and heat stress exposure lead to dissociation of the centromeres from nucleoli in a p38 MAPK kinase-dependent manner. This dissociation correlates with the alpha-satellite transcriptional response, and precedes heat stress-induced disruption of nucleoli protein nucleophosmin. Altogether, we provide evidence that inflammatory stresses induce changes in the chromatin landscape in human vascular endothelial cells by changing chromatin accessibility and decreasing the centromere-nucleoli association.

## DISCUSSION

Inflammation-induced changes at the different levels of chromatin organization are an under-investigated area of research yet are likely to be instrumental in determining the mechanisms of long-term cellular dysfunction and the cell fate switching, such as EndMT ^34^. Here, we focused on human vascular endothelial cells as a well-established model of inflammation-induced cellular dysfunction to investigate the effects of inflammatory stresses on chromatin accessibility and nucleoli-centromere dynamics.

Current knowledge regarding the chromatin changes in response to pro-inflammatory cytokines remains limited ^34^. One study revealed an intriguing phenomenon of a cell-type-specific prolonged inflammatory response that was accompanied by increased chromatin accessibility ^10^. The authors suggested that cells with prolonged high gene expression lacked adequate repression mechanisms, with potential changes in chromatin accessibility as one of the contributing factors. In another elegant work, the 3-7 days-long exposure to 5ng/ml of TNF alpha with high sugar did not lead to profound changes in Hi-C data but led to dramatic alterations in chromatin-bound RNA ^6^. The current work reports that short-term, six hours of cytokine exposure induced DARs that were much more local than megabases-long chromatin compartments in Hi-C data (Fig. 3). We also report an increase in chromatin accessibility related to EC dysfunction genes, ICAM1, VCAM1, EndMT and classic cytokine-inducible MAPK kinase and NFkB pathways.

The emerging data suggest that heat stress acts as a modifier of chromatin dynamics in human cells. For example, a dramatic rearrangement in chromatin organization during 1hr at 42ºC was reported in human embryonic stem cells ^35^, and the recent publication suggests significant changes in human K562 cells exposed to 42ºC at the chromatin accessibility level ^36^. Notably, these changes are reversible as the cells recover from stress. Our work agrees with previously reported changes in HS-induced chromatin accessibility, specifically DARs containing binding sites of heat shock transcription factors and a set of classic HS-response genes. Additionally, we report the DARs for EC-specific genes, including dysfunctional CAMs such as PECAM1 and anti-inflammatory proteins such as KLF4. We suggest that although the two inflammatory stresses, TNF-alpha and febrile-like HS, both induce EC dysfunction genes, these genes are rather stress-specific. The question regarding the complementary/cumulative effects of these stresses during inflammation requires further exploration.

The nucleolus is a cellular stress sensor in that during stress, it changes its function and protein composition, including chromatin modifiers, molecular chaperones, and nucleoli proteins such as NPM1 ^34,37–39^. Additionally, the nucleoli emerge as a gene silencing hub, including centromeric NADs in interphase cells, and thus, the possibility that stress-induced nucleoli changes affect nucleoli-associated chromatin domains arises. Here, we report that inflammatory stresses lead to the dissociation of centromeres from nucleoli. Centromere-nucleoli association in untreated interphase nuclei varies between cell types and healthy vs. cancer cells ^16^. We speculate that changes in dynamics during inflammation might have consequences such as an unscheduled gene activation of typically silenced chromatin domains. Our initial data support this notion as we observed an increase in pericentromeric transcripts upon stress and an “unscheduled” activation of the developmentally regulated gene ZNF423. Heat shock-induced transcription from pericentromeric regions was reported earlier in different human cells ^40–42^, as well as the elegant study pointing at the importance of nucleoli-centromere association for regulation of their transcriptional activity ^17^. Our data connected these findings in the context of inflammatory stresses in EC cells. We also observed dramatically decreased nuclear signal for NPM1 protein during cellular recovery from stress. This is an unexpected finding as a previous study suggested nucleoplasmic localization of NPM1 during short-term heat stress in human immortalized and tumor cells ^30^. This discrepancy may be attributed to the differences in cell types or temperature timing but indicate that the changes of NPM1 follow centromere-nucleoli dissociation rather than being a cause of a loss of centromeres-nucleoli contact. Thus, we suggest that inflammatory stresses change pericentromeric gene regulation in EC cells via p38MAPK but not via loss of nucleoli proteins NPM1 or nucleolin.

The limitation of this study is that it is unclear whether inflammatory stresses change histone modifications and whether these changes are reversible. It is also unclear whether an increase in centromere transcripts results in changes in heterochromatin features of centromere regions as well as NADs in general. Overall, we suggest that inflammatory stresses induce chromatin dynamics in EC cells in a stress-specific manner, and these changes are associated with transcriptional response, including cellular dysfunctional events.

## MATERIALS AND METHODS

*Cell culture and treatment*. Human aortic endothelial cells (HAECs; #PCS-100-011) and human umbilical vein endothelial cells (HUVECs; #PCS100010) were purchased from ATCC and cultured in endothelial cell growth basal medium-2 containing bullet kit growth factor supplements (EBM-2 [endothelial cell growth basal medium-2]; Lonza), 5% fetal bovine serum, 100 units/mL penicillin, 100 μg/mL streptomycin, and 2 mmol/L L-glutamine (Invitrogen).

For passaging, the old medium was aspirated, and cells were washed with 5 ml PBS per 10 cm plate. Then 1 ml of 0.05% (w/v) trypsin-EDTA was added, and the plate was incubated at 37°C until the cells detached. After, 5 ml RT medium was added per 10 cm plate to quench the trypsin, and the cells were transferred to a 15 ml conical tube and pelleted by centrifugation for 5 min at 200g at 4°C. Media was aspirated, and cells were resuspended in 5 ml of fresh room temperature warm medium. Cells were passaged at dilutions ranging from 1:2 to 1:10. Cultured human cells between passages 3 to 8 were used for experiments.

Cell treatment regimens are outlined in the table.

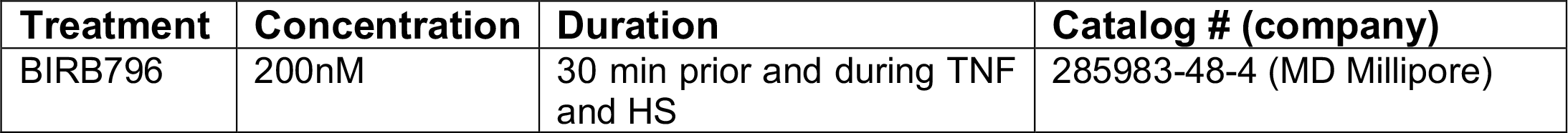
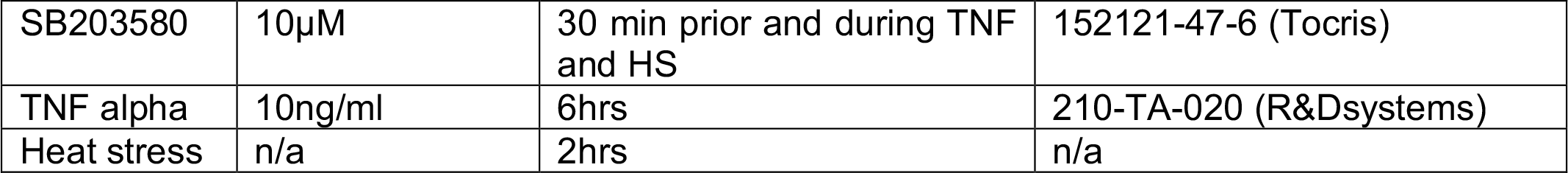

| Treatment | Concentration | Duration | Catalog # (company) |
| --- | --- | --- | --- |
| BIRB796 | 200nM | 30 min prior and during TNF and HS | 285983-48-4 (MD Millipore) |
| SB203580 | 10μM | 30 min prior and during TNF and HS | 152121-47-6 (Tocris) |
| TNF alpha | 10ng/ml | 6hrs | 210-TA-020 (R&Dsystems) |
| Heat stress | n/a | 2hrs | n/a |

*Quantitative PCR*. Control and treated HUVEC and HAEC cells were lysed with the TRIzol reagent (Life Science Technologies), and total RNA was extracted using the RNeasy Plus Micro Kit (Qiagen). Total RNA was reverse transcribed with oligo (dT) primers for cDNA synthesis using an iScript cDNA synthesis kit (Bio-Rad). The expression of mRNA was examined by quantitative PCR analysis using a QuantStudio 6 Flex Real-Time PCR System (Applied Biosystems). TaqMan assays were used to quantitate. The 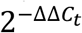 method was used for relative quantification of gene expression. Expression of *GAPDH* was used to normalize each sample.

| Probe Name | Number (Applied Biosystems) |
| --- | --- |
| ICAM1 | Hs00164932_m1 |
| IL6 | Hs00174131_m1 |
| PECAM1 | Hs00169777_m1 |
| VCAM1 | Hs01003372_m1 |
| KLF4 | Hs00358836_m1 |
| GAPDH | Hs99999905_m1 |

*RNA-seq*. HUVEC cells were grown in dishes, either left untreated or subjected to inflammatory stressor TNF alpha for 6hrs or febrile-mimicking heat stress for 2hrs and cryopreserved in 1ml of freezing media containing more than 0.4x10^6 cells. The experiments were done in biological triplicate. RNA-seq was done by Diagenode Inc. Specifically, nine cryopreserved samples were used for total RNA extraction using the Qiagen RNeasy mini kit (Qiagen Cat #74104). The RNA-seq experiment has been conducted by RNA-seq services (Diagenode Cat# G02030000). RNAs have been firstly quantified using Qubit™ RNA HS Assay Kit RNA HS Assay Kit (Thermo Fisher Scientific, Cat# Q32852). The RNA integrity was assessed using a Bioanalyzer, RNA 6000 Pico kit (Agilent Cat# 5067-1513,) on a 2100 Bioanalyzer system (Agilent). The samples with RINs above 7 were used for RNA-seq. Library preparation has been done with 250ng of input RNA using NEBNext Poly(A) mRNA Magnetic Isolation Module (NEB #E7490) followed by NEBNext Ultra II Directional RNA Library Prep Kit for Illumina (NEB #E7760) and NEBNext® Multiplex Oligos for Illumina® Index Primers Set 1 (NEB# E6440). The generated DNA libraries were purified using Agencourt® AMPure® XP (Beckman Coulter). Purified libraries were quantified, and their size was assessed with QIAxcel (Qiagen). The library pool was then sequenced on NovaSeq paired-end 50bp.

*ATAC-seq*. HUVEC cells were grown in dishes, either left untreated or subjected to inflammatory stressor TNF alpha for 6hrs or febrile-mimicking heat stress for 2hrs and cryopreserved in 1ml of media with DMSO containing 0.5-1.5x10^6 cells. The experiment was done in biological duplicate, paired with RNA-seq experiment. The ATAC-seq experiments have been conducted by the Diagenode ATAC-seq Profiling service (Diagenode Cat# G02060000).

Transposition. Cell viability and cell counting were evaluated by visualization under the microscope (AE2000; Matic). 200,000 cells were centrifuged at 500g before proceeding with the nuclei isolation and transposition reaction using the ATAC-seq Kit from Diagenode (Diagenode Cat #C01080002). According to the results of the nuclei isolation optimization, the Tween20/Igepal concentration in Lysis buffer 1 used was 0,1% with an incubation time of 3 minutes. 50,000 counted nuclei were transferred to a new tube and were centrifuged at 500g before proceeding with the transposition reaction. Isolated nuclei were lysed and transposed for 30 minutes at 37°C using the prokaryotic Tn5 transposase system (Nextera DNA library kit, Illumina, FC-121–1030). Transposed DNA was then purified on Diapure columns (Diagenode Cat# C03040001). Library preparation. Libraries were prepared from purified transposed DNA using NEBNext High-Fidelity PCR MasterMix (NEB, M0541) and Illumina indexing primers. The number of library amplification cycles was determined by qPCR analysis using NEBNext High-Fidelity PCR MasterMix (NEB, M0541) on LightCycler® 96 System (Roche). After their amplification, the libraries were size selected and purified using Agencourt® AMPure® XP (Beckman Coulter) and quantified using Qubit™ dsDNA HS Assay Kit (Thermo Fisher Scientific, Q32854). Finally, their fragment size was analyzed by High Sensitivity NGS Fragment Analysis Kit (DNF-474) on a Fragment Analyzer™ (Advanced Analytical Technologies, Inc.). Libraries were pooled and sequenced on an Illumina NovaSeq6000 in a 2 × 50-bp, paired-end mode.

### Bioinformatic analysis

#### RNA-seq data analysis

Quality of raw reads was assessed using FastQC (v0.11.9) (https://www.bioinformatics.babraham.ac.uk/projects/fastqc/). Given that the read quality was high and adaptor sequences were barely detected, no trimming was performed. Paired-end reads were mapped to the latest human reference genome T2T-CHM13v2.0 (Ensembl rapid release) using STAR (v2.7.10a) ^43^ in a two-pass mode, with the latest GTF (Ensembl rapid release, July, 2022) as gene annotation. A gene-by-sample count matrix was generated using featureCounts (v1.6.2) ^44^. All downstream statistical analyses were done using the R programming language (v4.1.0) Briefly, entries of genes with extremely low expression were first removed from the gene-by-sample count matrix. Differential gene analysis was performed using DESeq2 (v1.32.0) ^45^ with the batch effect considered. Genes with absolute values of log_2_(fold-change) ≥ 1 and BH method-adjusted *p* value ≤ 0.05 were considered as significantly differentially expressed genes (DEGs) ^46^. DEGs were visualized using the EnhancedVolcano package (v1.16.0) (https://bioconductor.org/packages/EnhancedVolcano/). Over-representation analysis of DEGs against KEGG pathways was performed using clusterProfiler (v4.0.5) ^47^. Gene set enrichment analysis of expressed genes in descending order of log_2_(fold-change) against the MSigDB get sets was carried out using the GSEAPreranked command of the GSEA software (v4.3.2) ^48^,^49^.

#### ATAC-seq data analysis

Quality of raw reads was assessed using FastQC (v0.11.9) (https://www.bioinformatics.babraham.ac.uk/projects/fastqc/). 3’-end adaptor sequences were trimmed using Timmomatic (v0.32) ^50^. The remaining paired-end reads were aligned to the latest human reference genome T2T-CHM13v2.0 (Ensembl rapid release) using bwa mem (v0.7.17) ^51^. Alignment files in the SAM format were first sorted by coordinates and converted into the BAM format using SAMtools (v1.16.1) ^52^. Subsequently, PCR duplicates were removed from the BAM files using the ‘MarkDuplicates’ command of the Picard tools (v2.27.5) (https://broadinstitute.github.io/picard/). The BAM files were further filtered to only keep concordant alignments with mapping quality ≥ 20 and insert sizes in the range of [38, 2000]. The resulting BAM files were name sorted using SAMtools. Post-alignment quality control was performed using the ATACseqQC package (v1.24.0) ^53^. Peaks per condition were called using Genrich (v0.6.1) with the BAM files of all biological replicates for a given condition and a q-value cutoff of 0.05. Consensus peaks-by-sample count matrix were generated using DiffBind (v3.4.11) ^54^. Transcription factor activity was inferred using the ChromVar package (v1.20.2) ^55^ and human core transcription factor motifs from the JASPAR database (v2016) ^56^. Peaks were associated to the nearest genes using the ChIPpeakAnno package (v3.34.1) ^57^. Differential peak analysis was conducted using DEseq2 (v1.32.0) ^45^. Peaks with absolute values of log_2_(shrunken fold-change) ≥1 and BH method adjusted *p* value ≤ 0.05, ^46^ were considered as significantly differential accessible regions (DARs). DARs were visualized using the Enhanced Volcano package (v1.16.0) (https://bioconductor.org/packages/EnhancedVolcano/). The nearest genes associated with the DARs were used for KEGG pathway overrepresentation analysis using clusterProfiler (v4.0.5) 5). Track views were gene-rated using the Integrative Genomics Viewer (v hg38) ^58^.

#### Immunocytochemistry and RNA-FISH

Single-molecule (sm) RNA-FISH experiments were performed according to the PixelBiotech Protocol (Schriesheim, Germany) and before immunocytochemistry. Briefly, cells were grown on sterile glass coverslips (Corning) until 60-80% confluency, treated with TNF alpha or febrile-like heat stress, triple washed with RT 1xPBS, and fixed with 4%PFA for 10 min at RT, washed with 1xPBS and stored in 70% ethanol overnight at 4ºC or used after an hour of incubation at 4ºC. Prior to hybridization, samples were washed twice with 2x SSC, 2M Urea (HuluWash) at RT and incubated with HuluProbe at 37ºC overnight. In order to eliminate the unbound probe, coverslips were washed four times with HuluWash at RT for 10 min, then with 1x PBS, and used for immunocytochemistry.

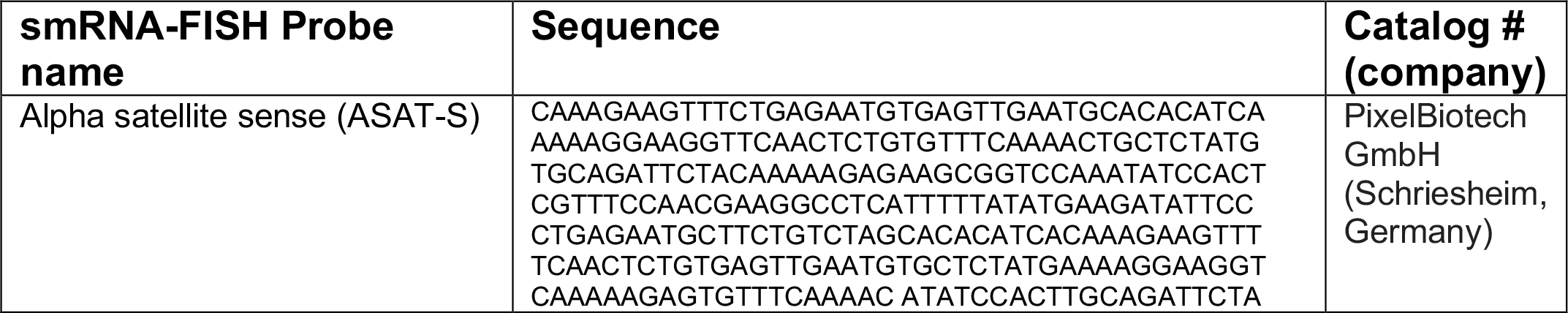
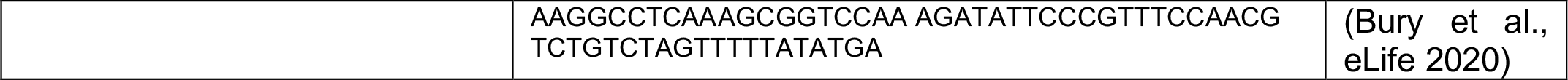

Immunocytochemistry as previously described. Briefly, cells were grown on sterile coverslips. After treatment, cells were rinsed with 1x PBS and blocked in 1x PBS/ 1% BSA for 15-30 min at RT. Primary antibodies were diluted in blocking solution and coverslips were incubated O/N at 4°C. Secondary antibodies for immunofluorescence were conjugated with: Alexa 488, Cy3 (Jackson ImmunoResearch, West Grove, PA). Samples were washed with PBS and incubated with secondary antibodies for 1 h at 37 °C. Coverslips were washed with PBS, 1x PBS / 0.1% TritonX-100, and 1x PBS at RT on a shaker, for 10 min each. Finally, the coverslips were rinsed twice with PBS and one time with water, and mounted using Prolong Gold antifade reagent (Invitrogen, cat # P36934).

| Antibody | Catalog #<br>(company) | Dilution | Fixation |
| --- | --- | --- | --- |
| nucleolin | ab22758 (Abcam) | 1:500 | MeOH;4%PFA |
| CREST | 15-235 (Antibodies Inc) | 1:100 | MeOH;4%PFA |
| nucleophosmin | (Abcam) | 1:500 | MeOH;4%PFA |
| pericentrin | ab4448 (Abcam) | 1:500 | MeOH |

#### Microscopy, software, and data analysis

Images were acquired with a Zeiss Axiovert 200M, a Perkin Elmer Ultraview spinning disc microscope, and a Hamamatsu ORCA-ER camera. (100x NA1.4 Plan-Apocromat Oil objective) and with Nikon A1RHD25 (100x NA1.45 Plan-Apocromat Oil objective). RNA-FISH probes and centromere/nucleoli association were counted through Z-stacks manually and were scored as “associated” when no gap between the nucleolar marker and the centromere marker was detected throughout the Z-stack. For images, fluorescence range intensity was adjusted identically for each series of panels. *Z* stacks are shown as 2D maximum projections or as a single optical section (MetaMorph, Molecular Devices, or Nikon Elements AR software). All images across each experimental series were taken using the same microscope setting (such as laser power and gain) to allow equal comparison of fluorescence levels of samples. All statistical analysis was done using GraphPad Prism software.

## AUTHORS CONTRIBUTIONS

A.C., H.L. (Heather Learnard) made experimental contributions; M.K., S.K., and J.K. provided invaluable input in endothelial cells biology and manuscript writing; H.L. (Haibo Liu) and L.Z. did bioinformatic analysis and contributed to writing; A.V. developed the concept, made experiments, and wrote the manuscript, and all authors critically read and commented on the manuscript.

## ACKNOWLEDGEMENTS

This work was supported by the American Heart Association Career Development Award to A.V. (grant number 856074), Brigham and Women’s Hospital Heart and Vascular Center Junior Faculty Award to S.K., NIH 1R03TR004452-01 to S.K. and NIH 5R01HL151626 to J.F.K. The authors would like to thank Paul Kaufman for support; Gagandeep Kaur, Serena S. David, and Natalia Naumova for their support and critical reading, and apologize to all scientists whose work was not cited due to space limitations.

## DECLARATION OF INTEREST

The authors declare that the research was conducted in the absence of any commercial or financial relationships that could be construed as a potential conflict of interest.

## Notes

### Competing Interest Statement

The authors have declared no competing interest.

