## Supplemental Figure 1 for "Inflammatory stress-mediated chromatin changes underlie dysfunction in endothelial cells"

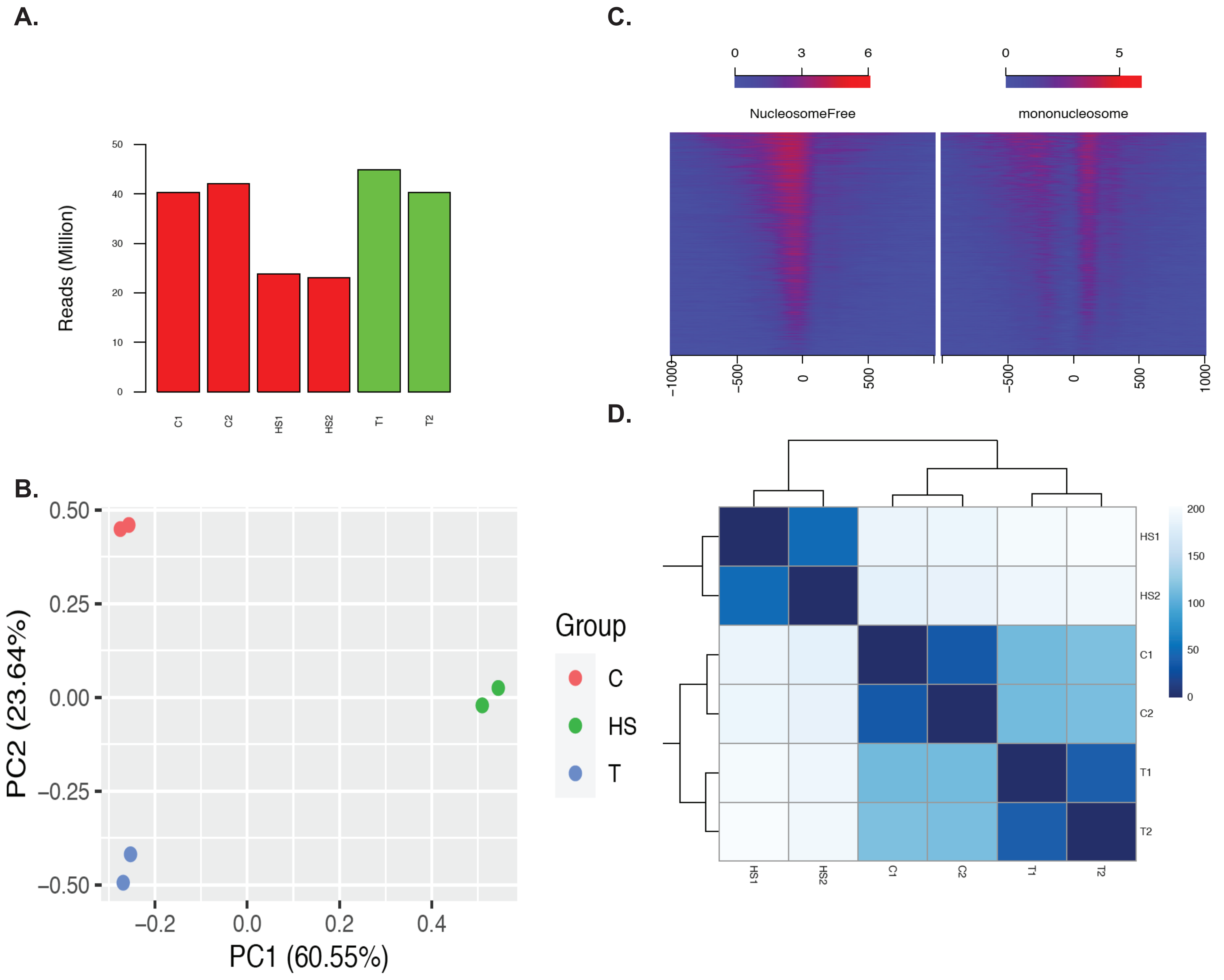


Figure S1. Related to Fig.1. ATAC-seq on HUVEC cells. A. Number of reads assigned to consensus peaks. B.Principal component analysis. C. Control ATAC-seq TSS signal read heatmap. D. Heatmap showing original samples distance.
