## Supplemental Figure 2 for "Inflammatory stress-mediated chromatin changes underlie dysfunction in endothelial cells"

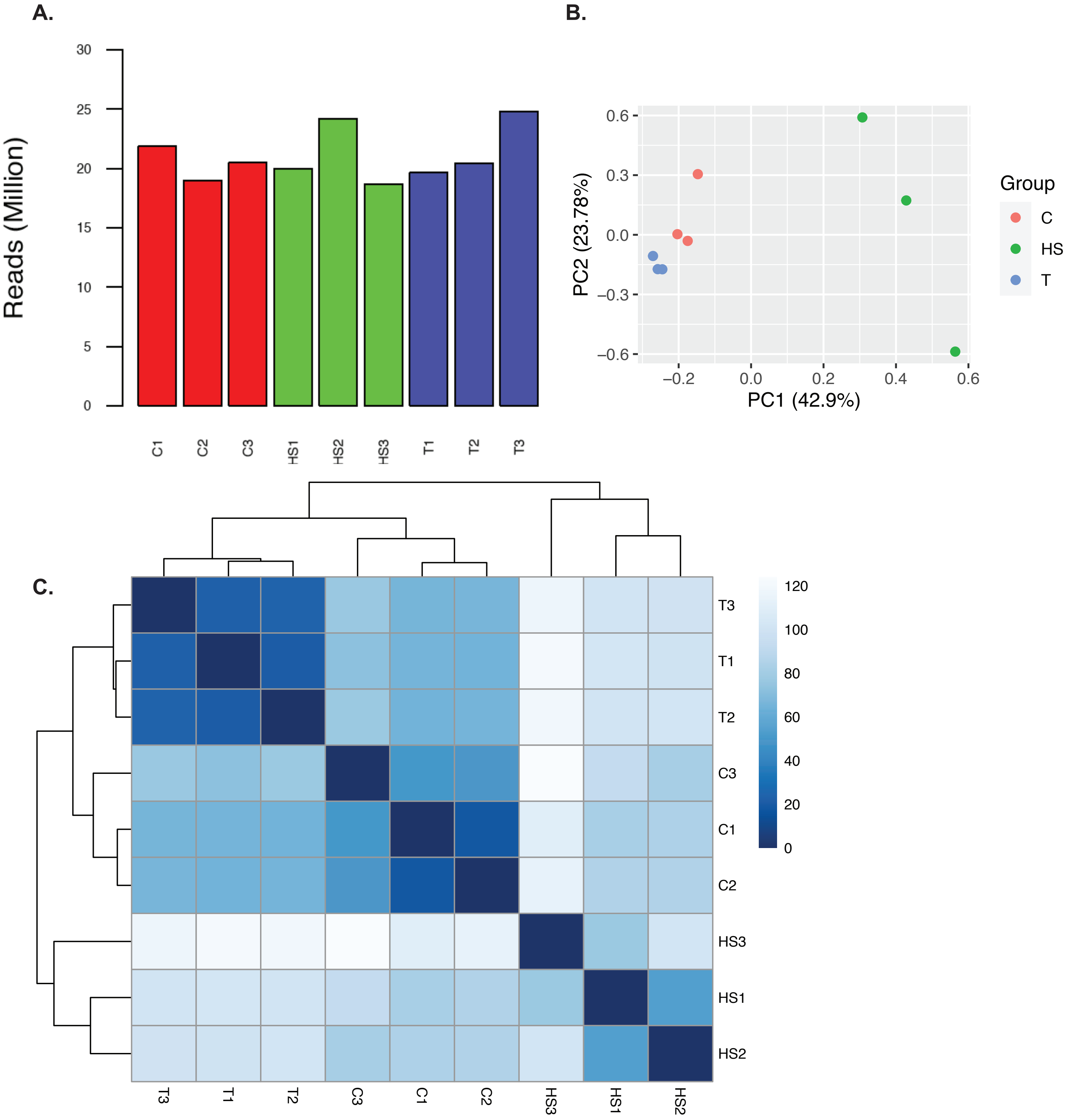


Figure S2. Related to Fig.2. RNA-seq on HUVEC cells. A. Number of reads of the three biological triplicates, where C1, C2, and C3 are control samples from distinct biological replicates; HS1, HS2, HS3 are heat shock samples, and T1, T2, and T3 are samples treated with TNF-alpha. B.Principal component analysis of the three biological replicates. C. Heatmap showing original samples distance.
