## Supplemental Figure 3 for "Inflammatory stress-mediated chromatin changes underlie dysfunction in endothelial cells"

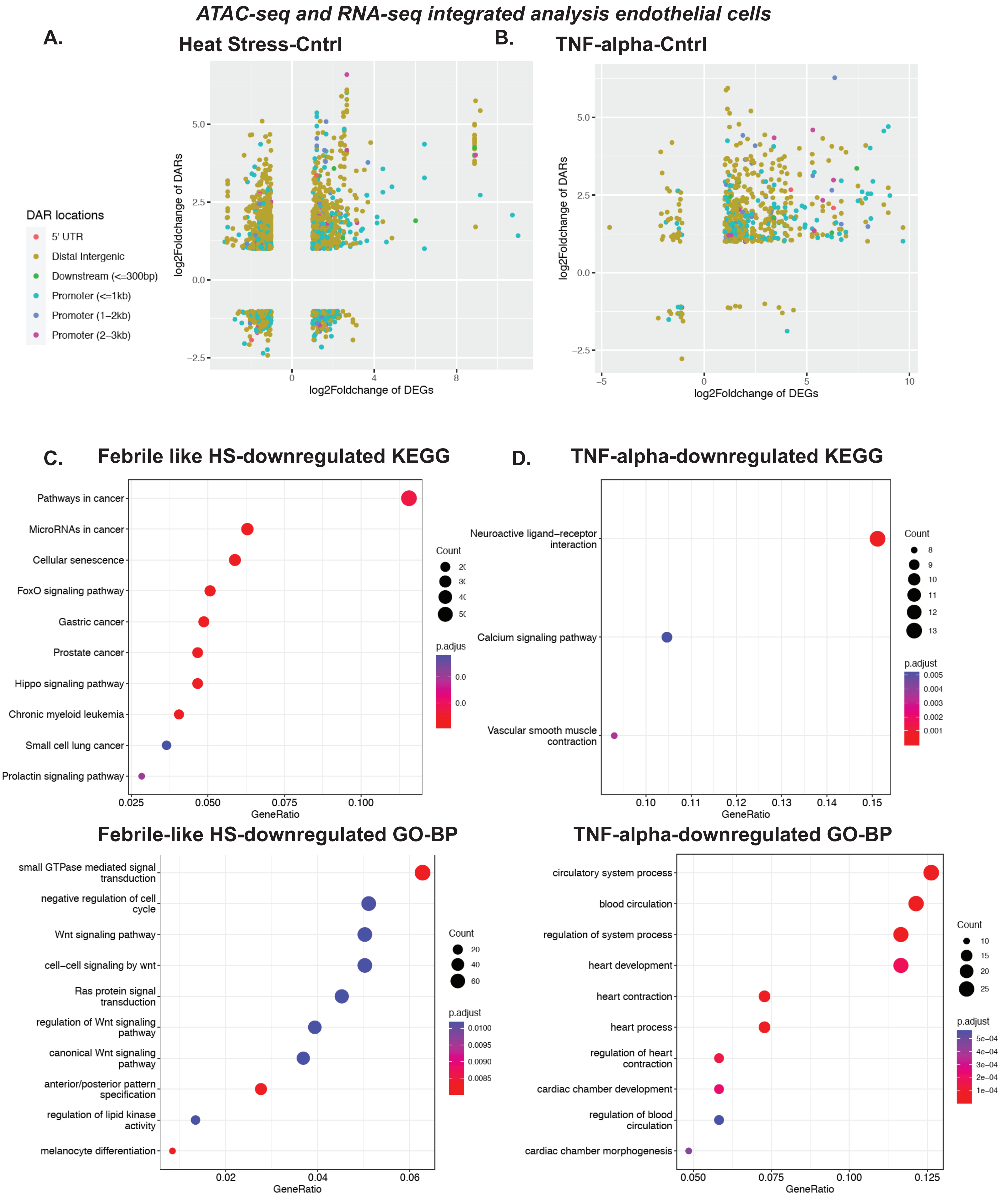


Figure S3. Related to Fig.3. ATAC-seq and RNA-seq integrative analysis. A. Genomic features of ATAC-seq and RNA seq in HS-C samples. B. Same as A but in T-C samples. C. KEGG and GO-BP analysis of downregulated pathways in HS-C samples. D. Same as C but in T-C samples.
